# eCOMET: An R package for evaluating metabolic diversity and enrichment from LC-MS/MS data to test ecological hypotheses from individuals to ecosystems

**DOI:** 10.64898/2026.06.02.729701

**Authors:** Min-Soo Choi, Dale L. Forrister, Guillaume J. Dury, Kyo Bin Kang, Brian E. Sedio, Youngsung Joo

## Abstract

- Methods in metabolomics have grown exponentially in recent years, providing new insight into the ecological function and evolutionary impact of diverse plant metabolites. Metabolomics requires a command of numerous tools, the outputs of which are typically integrated through in-house, custom code that presents a workflow bottleneck and a barrier to entry for researchers in ecology, evolution, and behavior who may benefit from adding a metabolomics perspective to their research.
- We introduce *eCOMET*, an R package for integrating and harmonizing the outputs of common metabolomics bioinformatics tools and conducting statistical analyses and data visualization methods useful for ecological metabolomics.
- Our package combines metabolome feature metadata with quantification tables (e.g., *mzmine*), feature dissimilarity matrices (e.g., modified cosine and *DreaMS*), and feature annotations (e.g., *SIRIUS*) into a cohesive R data object to facilitate downstream analyses, including the calculation of diversity and disparity metrics and differential accumulation analysis.
- Our goal is to make metabolomics accessible to a wider range of researchers in ecology, evolution, and behavior to unlock the potential of ecological metabolomics to generate novel insight in these fields.

## Introduction

Metabolites constitute a key functional language of ecosystems, serving as the chemical interface that shapes biological communities and drives ecosystem processes. While chemical ecology has successfully identified the ecological functions of diverse metabolites, the field is rapidly expanding toward a holistic, metabolome-scale perspective (Endara *et al*., 2023). This transition to metabolomics is facilitated by advances in analytical chemistry, especially liquid chromatography-tandem mass spectrometry (LC-MS/MS) (Sedio, 2017; Li & Gaquerel, 2021). Modern mass spectrometry methodologies now enable sub-ppm sensitivity and rapid scan speeds for acquisition of tens of thousands of tandem mass spectra per run (Defossez *et al*., 2023). These technological advances allow researchers to move beyond well-known metabolites and explore the wider chemical space observed in a diverse set of organisms across a variety of ecological and evolutionary contexts (Defossez *et al*., 2021; Forrister *et al*., 2022; Sun *et al*., 2024). Collectively, the field in which researchers apply untargeted mass spectrometry-based metabolomics to study the chemical phenotypes of organisms in their ecological and evolutionary contexts (Walker *et al*., 2023) can be referred to as *ecological metabolomics (hereafter, ecometabolomics)*.

Extracting biological meaning from untargeted metabolomics data requires chaining together several distinct analytical steps (Schmid *et al*., 2023; Heuckeroth *et al*., 2024). Raw LC-MS/MS data are first processed into feature tables through peak detection, chromatographic alignment, and quantification, producing feature definitions and their abundances across samples. Numerous platforms are available to facilitate these essential steps, including *mzmine* (Heuckeroth *et al*., 2024), *MS-DIAL* (Tsugawa *et al*., 2019), *Asari* (Li *et al*., 2023), and *XCMS* (Smith *et al*., 2006). The outputs of these tools are typically feature quantification matrices and molecular networks, which are then fed into the next step, which attempts to annotate identified features with putative structural or chemical-class information. Lastly, feature tables, molecular networks and annotation outputs must be integrated into a variety of statistical analysis to identify patterns in feature abundance across sample groups. Each of the aforementioned steps is underpinned by its own ecosystem of software, and no single program spans the full workflow. As a result, practitioners must move data between multiple, independently developed tools and typically write custom scripts to reconcile their formats before any interpretation can begin. This often-ignored integration step acts as a barrier to entry, because it is not well documented by existing tools, and thus requires learning to navigate a complex ecosystem of tools. Moreover, integration code is rarely standardized across lab groups or studies, thus hindering reproducible science.

The number of experimental mass spectra has grown exponentially in the past 5 years (Bushuiev *et al*., 2025), yet fewer than 10% of detected molecular features match standards of known compounds through spectral library matching (Aron *et al*., 2020). In response to this annotation gap, the annotation step has seen by far the greatest proliferation of new methods, with the development of many innovative tools that provide multi-layered information that supplements the unidentified spectra. MS/MS spectral similarity networking revolutionized the way of handling unannotated spectra, by suggesting possible structural similarities (Watrous *et al*., 2012; Wang *et al*., 2016). Machine-learning frameworks such as *MS2DeepScore* (de Jonge *et al*., 2026), *Spec2Vec* (Huber *et al*., 2021), and *DreaMS* (Bushuiev *et al*., 2025) can generate more reliable embeddings to assess structural similarity of features across entire datasets, complementing the modified cosine similarity, the spectral similarity metric used for the first developed MS/MS networking. *In silico* annotators, such as *CSI:FingerID* (Dührkop *et al*., 2015), *MolDiscovery* (Cao *et al*., 2021)*, COSMIC* (Hoffmann *et al*., 2022) and *ICEBERG* (Wang *et al*., 2026a) match the unidentified spectra against structural libraries and rank the possibilities by probability. *CANOPUS* (Dührkop *et al*., 2021) offers hierarchical chemical classifications, and *MS2LDA* (van der Hooft *et al*., 2016) provides substructural motifs (Beniddir *et al*., 2021). More recently, advances in machine learning introduced *de novo* annotation tools, such as *MSNovelist* (Stravs *et al*., 2022) and *MetGenX* (Wang *et al*., 2026b). Often combined, these approaches are particularly valuable in ecological research, where the study subjects are often non-model organisms, the metabolites of which are therefore especially poorly represented in existing spectral repositories (Uthe *et al*., 2021).

The final step, statistical analysis and biological interpretation, draws on all of the preceding outputs—feature tables, molecular networks, and annotations—and is again served by its own set of dedicated platforms. *MetaboAnalyst* (Pang *et al*., 2024) provides a broad, graphical user interface-based statistical workbench oriented toward clinical and biomedical study designs. *FBMN-Stats* (Pakkir Shah *et al*., 2025) offers a protocol and companion code for statistical analysis of GNPS feature-based molecular networking output, and *FERMO* (Zdouc *et al*., 2025) supports hypothesis-driven prioritization of molecular features, particularly in natural product discovery. Across these platforms, the dominant objectives are ranking features by statistically significant abundance patterns and characterizing samples by their annotated constituents.

However, ecological metabolomics poses analytical needs these existing tools and platforms were not designed for. Ecological hypotheses are often framed around the chemical diversity and dissimilarity across samples, referred to as chemical alpha and beta diversity, respectively (Sedio, 2017; Wetzel & Whitehead, 2020; Thon *et al*., 2024). Since multiple species often share few compounds but contain many structurally related metabolites, leveraging chemical similarity between compounds enables a more chemically relevant way of diversity calculation (Petrén *et al*., 2023). This approach is analogous to the concept of phylogenetic diversity, where the relatedness between species is introduced to account for the phylogenetic relationships among species when calculating ecological diversity (Faith, 1992; Lozupone & Knight, 2005).The low annotation rate of metabolomics data can be compensated by utilizing spectral similarity (Tripathi *et al*., 2021), because structurally related compounds coincide in similar biological contexts and the structural relatedness can be inferred from mass spectra. While several previous approaches required chemical structure to calculate structural similarity (Junker, 2018; Petrén *et al*., 2023), employing spectral similarity enabled chemical diversity calculation even without exact structural annotations (Sedio *et al*., 2017). Leveraging these diverse bioinformatic methods requires a command of numerous tools, the outputs of which are typically integrated through in-house, custom code that presents a workflow bottleneck and a barrier to entry for researchers in ecology, evolution, and behavior who may benefit from adding a metabolomics perspective to their research.

We introduce *eCOMET*, an *R* package for ecological metabolomics that pursues two complementary goals: First, to integrate outputs from common upstream metabolomics tools—*mzmine* (Heuckeroth *et al*., 2024), *SIRIUS* (Dührkop *et al*., 2019), and *GNPS* (Aron *et al*., 2020) to name a few—into standardized, goal-oriented analytical workflows tailored to the needs of ecometabolomics. Second, to lower the barrier to entry for ecologists to use metabolomics data by making the analytical assumptions, limitations, and decision points of each step explicit and navigable. By organizing core functionalities such as chemically informed diversity and dissimilarity quantification and differential accumulation analysis, *eCOMET* facilitates data exploration through an ecological lens and simplifies the testing of ecological hypotheses using untargeted metabolomics datasets. Below, we describe the structure of the *eCOMET* package and demonstrate its utility through two case studies.

## Methods

### Conceptual overview of the package eCOMET and its workflows

*eCOMET* aims to manage the inherently heterogeneous ecological and metabolomic datasets by first integrating common metabolomics datasets generated through various workflows into a single object, and then by using the object and ecological (as well as evolutionary or behavioral) datasets for downstream analyses (Fig. 1). At the data integration stage (Fig. 1a and Fig. 2), processed LC-MS data are integrated into a mass-spectrometry metabolomics object (*mmo*), which serves as the analytical hub for statistical analyses and visualization methods available in the package. The unified structure ensures data integrity is maintained. Once data are integrated, the *mmo* enables streamlined imputation, normalization, and filtering (Fig. 1b). After the data are adequately prepared, the package offers several options designed to cover a wide range of needs from ecology and evolutionary biology (Fig. 1c–f).

**Figure 1.**
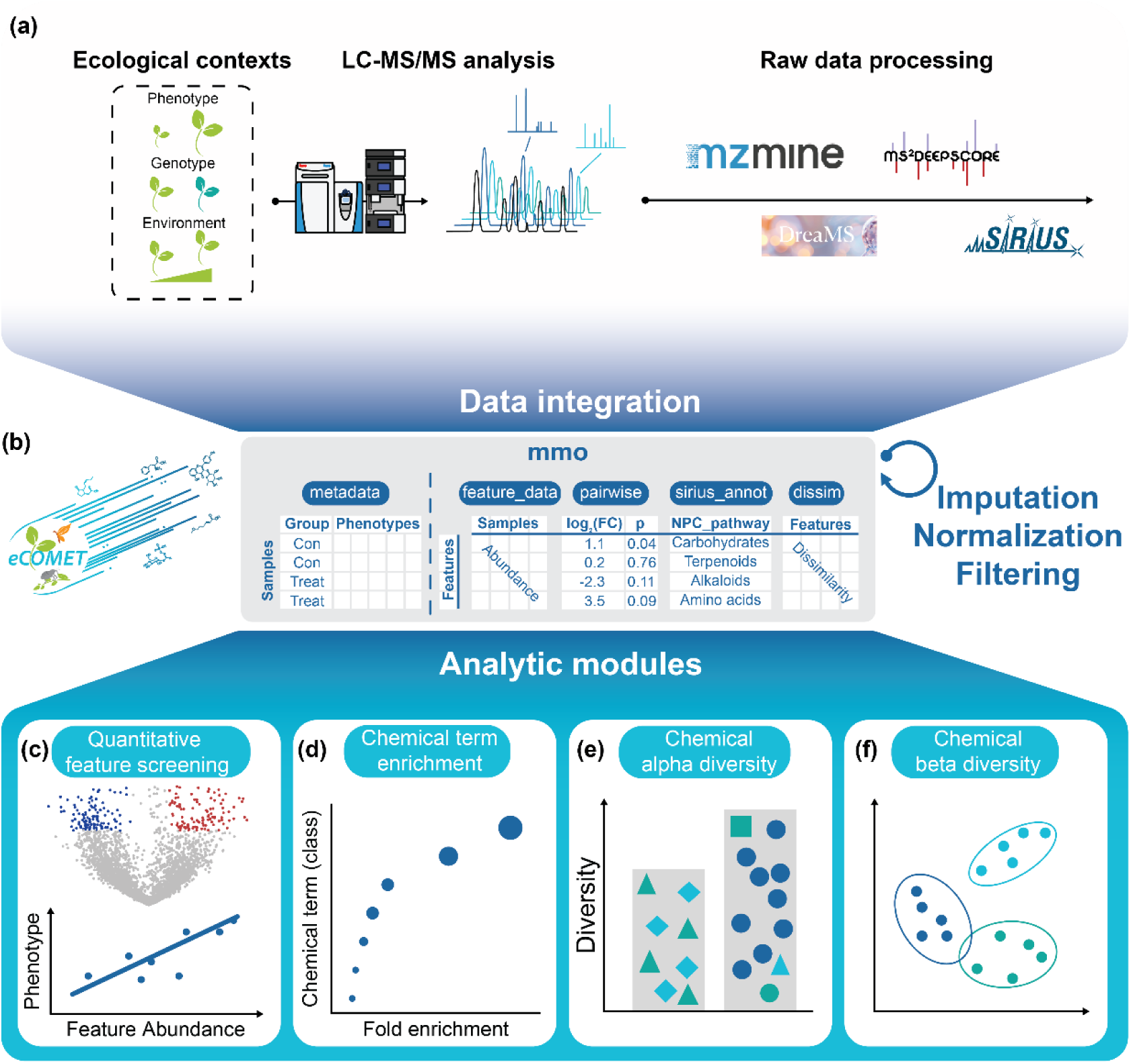
Overview of eCOMET workflows. (a) Samples from various ecological contexts are targeted for metabolomic analyses, which of the raw data files are processed through various tools including mzMine, SIRIUS, DreaMS, and MS2DeepScore. (b) The inputs are integrated into a central object mmo for imputation, normalization, and filtering for further analyses. (c-f) Analytic modules loads inputs from mmo to perform (c) differential accumulation analysis and correlation analysis, (d) chemical term enrichment analysis, (e) chemical alpha diversity quantification, and (f) chemical beta diversity quantification.

**Figure 2.**
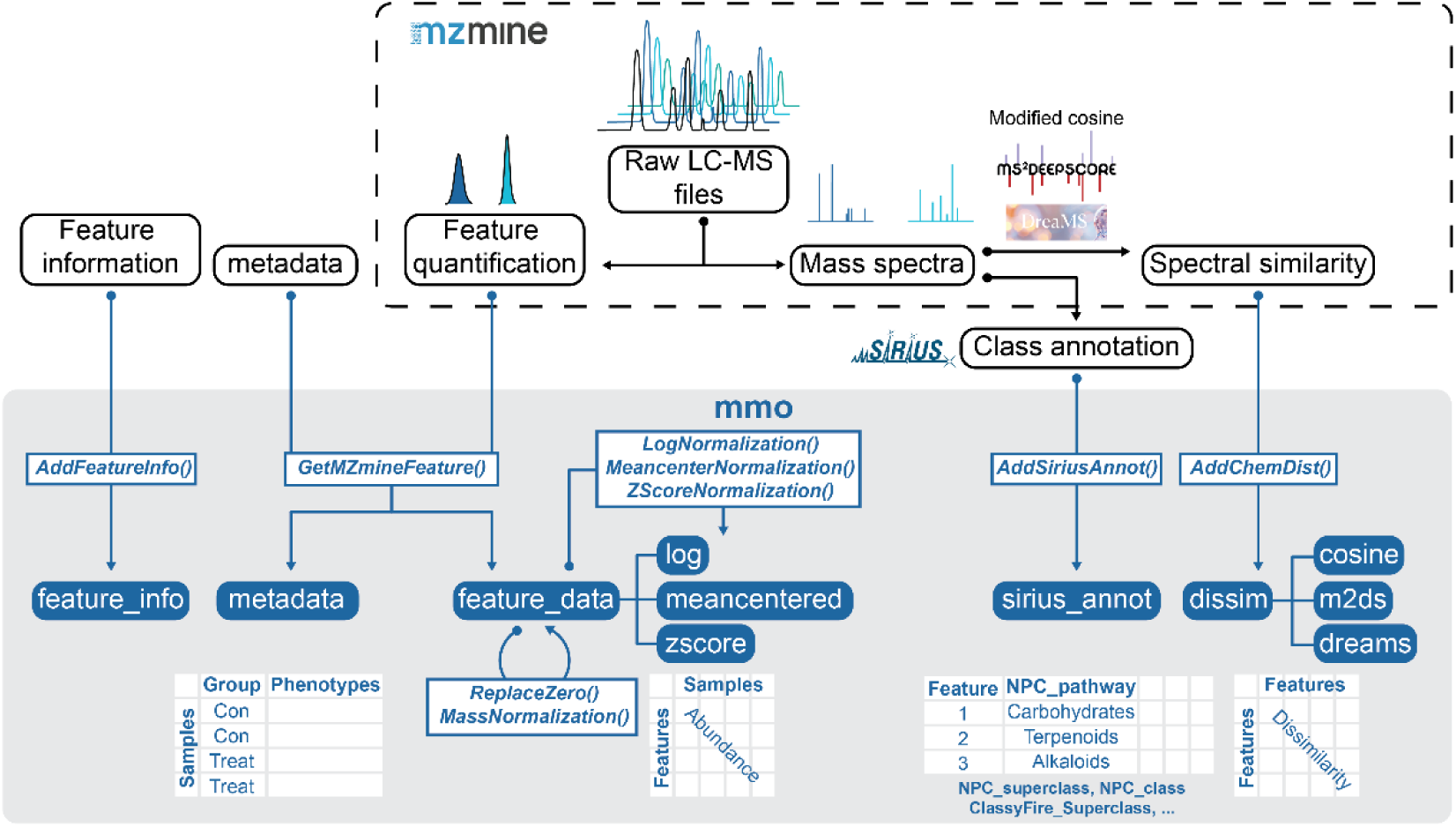
Data integration module of eCOMET. An mmo object is first generated by the function GetMZmineFeature(), then multiple importing functions add inputs to the mmo. The feature quantification matrix is normalized in various methods, with the feature IDs are paired with class annotation information and spectral dissimilarity matrix. The metadata provides any biologically relevant data for downstream analyses.

### Data inputs

We assume that an ecometabolomic study will contain two, and commonly four, types of data:

- Sample metadata: one or more quantitative or categorical variables defined at the level of individual samples that describe individual phenotype, population, experimental treatment, species identity, or taxonomic class
- Sample × feature quantification table: an aligned feature abundance table across samples that can be used for uni– and multivariate statistics
- (Optional) Feature annotations and definitions: mass-to-charge ratios, retention time and annotations consisting of predicted structures, classifications or chemical properties
- (Optional) Feature dissimilarity matrices: pairwise similarity matrices calculated from diverse spectral similarity metrics (e.g., modified cosine, *MS2DeepScore*, and *DreaMS*)

While a variety of software and pipelines can generate these types of data, *eCOMET* integrates them into a standardized object that can be transformed, filtered, statistically analyzed, and visualized.

### Data integration and construction of the mmo object

*eCOMET* takes four types of preprocessed files as input: 1) a feature-by-sample table produced by *mzmine* (version > 4), 2) a feature-by-feature similarity table produced by molecular networking in *mzmine*, 3) a class-level annotation table of features produced by *SIRIUS*, and 4) a metadata table describing the grouping of samples, including by phenotypes or by environmental variables. More detailed explanations, examples, and guides to run *eCOMET* are available in the supplementary materials and GitHub.

To integrate the input files into an *mmo* object, the following functions are used sequentially: The function *GetMZmineFeature* takes an *mzmine* feature-by-sample quantification table and its associated metadata to construct an *mmo* object containing *feature_data* and *metadata*. Even with the basic gap-filling module of *mzmine*, the *feature_data* contains empty or zero values which can impair downstream analyses including PCA and differential accumulation analysis. To avoid this, empty values can be replaced with imputed values of 1 or half of the minimum non-zero value in the row using the function *ReplaceZero*. Either the raw feature abundance data or the imputed data can be selected depending on statistical needs. For example, pairwise comparison should generally be performed on imputed data, whereas richness evaluation should be performed using the original feature table.

### Feature Abundance Normalization and Transformations

The function *MassNormalization* normalizes the raw peak area values by sample mass if this information is provided in the metadata. Since normalizations of the peak area values are often needed for downstream analyses, functions are provided to log normalize (*LogNormalization*), mean-center (*MeancenterNormalization*), and Z-transform (*Znormalization*) values of the *feature_data*.

### Feature Annotation and Feature-by-Feature Pairwise (Dis) similarity

The function *AddSiriusAnnot* integrates the outputs of SIRIUS, including predicted structures and classifications, into the *mmo* object as *sirius_annot*. Feature-by-feature similarity metrics are transformed into dissimilarity and added to the *mmo* object by the function *AddChemDist*. These include cosine (*cos.dissim*), *MS2DeepScore* (*m2ds.dissim*), and *DreaMS* embeddings (*dreams.dissim*). The final structure of the *mmo* is illustrated in Fig. 2.

### Filtering of the mmo Object

To subset the *mmo* based on sample groups, or a list of specific samples or features, the function *filter_mmo* can be used to create a new *mmo* object containing a subset of the original *mmo* object. The function preserves the integrity of the *mmo* by filtering using metadata, feature abundance, feature definitions, or annotations. To facilitate easy filtering during visualization and analysis the *filter_mmo* function can be called within most *eCOMET* functions, using the same logic and provided parameters for sample, group, or feature.

### Scalability and streamlining of statistics and visualization

Ecological metabolomics studies rely on pre-existing R packages to perform various statistical tests required for many common research objectives. The goal of *eCOMET* is not to re-invent existing analyses, but rather to provide a standardized way to integrate pre-processed metabolomics data into a suitable format, the *mmo*, for commonly performed statistical analyses in ecology. To streamline the integration of upstream preprocessed metabolomics data and downstream statistical analysis and visualization we built all functions around the standardized *mmo* structure. Additionally, we provide core functionality for users to specify parameters such as normalization, to define the quantification data, or filter a subset of features, groups or samples.

Below, we outline the analysis methods currently implemented, explain how they are used in ecometabolomics studies and provide references for the underlying packages. Most statistical functions in *eCOMET* provide relevant plots for rapid visualization and manual inspection of the results. Where appropriate, functions also provide printed statistical results and a data frame with the underlying data used to generate visualizations. The output can therefore be used for further analysis, alternative visualization, or sharing, thereby improving scalability and reproducibility.

### Differential accumulation (DA) analysis

Differential expression analysis is a fundamental workflow in transcriptomics, in which the mRNA read counts of genes are compared across groups to identify those whose expression differs systematically with treatment or condition. The same statistical logic can be applied to untargeted metabolomic data, where peak areas of molecular features replace read counts of genes as the quantitative units of comparison. We refer to this approach as differential accumulation (DA) analysis and implemented it in eCOMET through the following functions: *PairwiseComp* takes the *mmo* object and names of two groups (each being the numerator and the denominator) to calculate log_2_ fold change, *p*-value from Welch t-test, and Benjamini–Hochberg (BH) adjusted *p*-value of each feature in pairwise group comparisons. The results are stored in *mmo$pairwise* as two additional columns each time *PairwiseComp* is called. The results of DA analyses can be visualized in a volcano plot by the function *VolcanoPlot*. The differentially accumulated molecular features (DAMs) can be extracted by the function *GetDAMs* by user-set thresholds of p-values and the fold changes.

### Phenotype association analysis

While DA analysis identifies features that differ between discrete groups, many ecological and physiological studies instead seek to relate metabolite abundance to continuous traits or phenotypes, such as growth rate, herbivory, or environmental gradients (Wu *et al*., 2018). *eCOMET* implements such association analyses through the following functions. The association analysis between certain feature abundance and quantitative phenotype of interest which can be provided by the metadata is implemented in the function *FeaturePhenotypeCorrelation*. Screening such association along all features can be performed by *ScreenFeaturePhenotypeCorrelation*. The association can be calculated by one of following: linear model, linear mixed model with random effects, Pearson correlation, Spearman correlation, and Kendall correlation.

### Chemical class enrichment analysis

Comparative and correlative analyses inomics studies typically produce long lists of genes, proteins, or metabolites that differ across conditions, but interpreting these lists biologically requires aggregating individual features into functionally meaningful groups. Overrepresentation analysis (ORA) addresses this need by testing whether predefined functional classes are enriched in a given feature list relative to the background (Wieder *et al*., 2021). eCOMET extends this logic to chemical classification through two complementary approaches: chemical class overrepresentation analysis (cORA) for discrete feature sets, and metabolite set enrichment analysis (MSEA) for continuously ranked features. To conduct cORA, the function *CanopusLevelEnrichmentAnal* calculates overrepresentation of certain chemical class in given set of features. The function first takes inputs of *mmo*, a list of features, and the level of class annotation (from CANOPUS) to be tested. Class ontology from either ClassyFire (Djoumbou Feunang *et al*., 2016) or NPClassifier (Kim *et al*., 2021) can be used (one of ‘NPC_pathway’, ‘NPC_superclass’, ‘NPC_class’, ‘ClassyFire_superclass’, ‘ClassyFire_class’, ‘ClassyFire_subclass’, ‘ClassyFire_level5’, or ‘ClassyFire_most_specific’). The abundance of each chemical class in total features list and the list provided are calculated, then the Fisher’s exact test is used to calculate the significance of overrepresentation and BH method is applied to correct for multiple testing. The result of cORA, comprising the name of chemical classes, (adjusted) p-values, counts in the provided list and total feature list, and fold enrichment, can be used for further visualizations. The functions *CanopusListEnrichmentPlot* and *CanopusLevelEnrichmentPlot* provide a preset visualization of the cORA results.

Complementary to the cORA, which requires a list of features, a metabolite set enrichment analysis (MSEA) is also implemented to identify the chemical classes which are associated with given continuous values for each feature. An *mmo* object, the level of chemical class to be investigated, names and scores of features to be analyzed are used as inputs of the function *MSEA* then transferred to fgsea (Korotkevich *et al*., 2021), to calculate normalized enrichment scores, p-values and adjusted p-values for each chemical class and generate a preset plot.

### Chemical diversity analyses

In ecology, alpha diversity is the mean species diversity in a site at a local scale. When applied to metabolomics, chemical alpha diversity refers to the diversity of molecular features within a sample. A popular alpha diversity index is the Hill number, which considers both abundance and richness, it corresponds to the effective number of features—the number of equally-abundant features needed to obtain the same mean proportional feature abundance as that observed in the sample (Hill, 1973; Chao *et al*., 2014). The Hill number framework has been adapted to metabolomic data in the R package *ChemoDiv* (Petrén *et al*., 2023). In addition to the classical Hill number, the “functional Hill numbers” are calculated by weighing the contribution of each compound by its structural dissimilarity to all other compounds in the dataset. All else equal, this approach increases the diversity value of samples containing more structurally distinct compounds. If given structures of compounds, the package *ChemoDiv,* calculates the similarity between compounds to calculate functional Hill numbers. However, this limits the calculation to only features with structural annotations, often a small subset of features, and structural prediction is typically the least confident annotation available, compared to molecular formula or chemical class predictions. Therefore, we implemented the functional Hill numbers calculation by directly providing spectral similarity matrix (e.g., modified cosine or DreaMS embeddings similarity). The function *GetAlphaDiversity* calculates the Hill number (chemically naïve) or functional Hill number (chemically informed), by user-specified *q* values (where *q* is the sensitivity to rare vs. abundant features, with higher numbers corresponding to higher sensitivity to abundant features). For more detailed information on the functional Hill number, refer to (Chao *et al*., 2014; Petrén *et al*., 2023).

Chemical beta diversity is defined by the distance (or dissimilarity) between samples. While classical methods to calculate sample-to-sample distance are to implement Bray–Curtis or Jaccard dissimilarity, chemically informed analogs that leverage the spectral similarity between features include CSCS (Sedio *et al*., 2017) and UniFrac dissimilarity (Junker, 2018). The function *GetBetaDiversity* transforms the feature abundance table into a relative proportion table, then calculates one of Bray–Curtis, Jaccard, CSCS, or UniFrac dissimilarities between all pairs of samples. When either UniFrac or CSCS is selected, the spectral similarity metric needs to be chosen among cosine, MS2DeepScore (de Jonge *et al*., 2026), and DreaMS embeddings. Also, various alpha values for UniFrac from unweighted to fully weighted are allowed.

### Multivariate analyses

Multivariate analyses such as Principal Component Analysis (PCA), Partial Least Squares-Discriminant Analysis (PLS-DA), Non-metric Multidimensional Scaling (NMDS), and Principal Coordinates Analysis (PCoA) are implemented in eCOMET along with appropriate 2-dimensional visualization methods. The functions PCAplot and PLSDAplot require only the *mmo* object, while the functions *NMDSplot* and *PcoAplot* require an additional beta-diversity matrix, which is a distance matrix of samples output by the function *GetBetaDiversity.* The function *adonis2* from *vegan*package and *pairwiseadonis* (https://github.com/pmartinezarbizu/pairwiseadonis) is used to conduct PERMANOVA subsequent to multivariate analyses and the results are provided together with the calculated coordinates.

## Results and Discussion

To ensure the package’s comprehensiveness, we surveyed foundational questions in ecometabolomics, mapping each to its required statistical frameworks and necessary data structures (Table S1). We observed that while the ecological contexts vary, studies often share underlying analytical requirements. These include comparing metabolite abundances across groups, correlating environmental variables with chemical profiles, or quantifying chemical dissimilarity across communities. After identifying these commonalities, we organized the package into four core analytical modules capable of addressing the majority of current eco-metabolomic hypotheses (Fig. 1c-f).

**Module 1. Quantitative feature screening** The first module identifies molecular features that drive ecological patterns (Fig. 1c and 3a), finding associations with categorical treatments, or continuous covariates. Differential Accumulation (DA) analysis compares feature abundances across discrete sample groups to identify Differentially Accumulated Metabolites (DAMs), which are visualized via volcano plots and Venn diagrams. Complementary to this, correlation analyses are used to identify features whose abundances are associated with continuous or discrete phenotypic traits or environmental variables (e.g., herbivore performance, soil nutrient gradients) for hypothesis generation.

**Figure 3.**
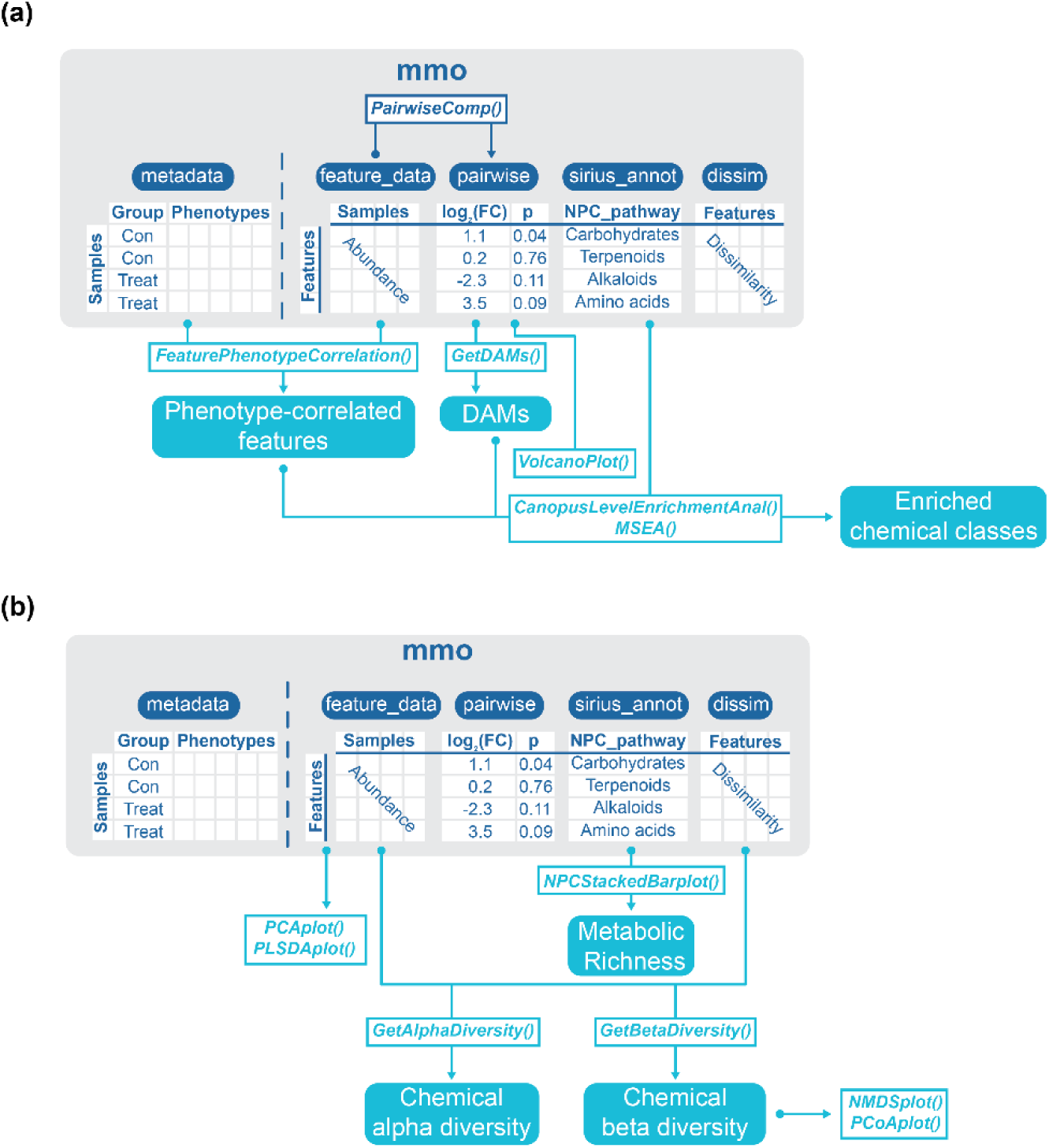
Analytic modules of eCOMET. (a) The feature quantification data and metadata are used to perform differential accumulation analysis and correlation analysis to generate lists of features. The features then can be further used for chemical term enrichment analysis. (b) PCA or PLS-DA plots can be generated solely by the feature quantification matrix. The spectral dissimilarity matrix and feature quantification matrix is processed to quantify chemical alpha and beta diversities. The class annotations can be visualized as metabolic richness.

**Module 2. Chemical term enrichment analyses** A primary bottleneck in ecometabolomics is that the majority of DAMs or phenotype-associated features remain unannotated based on spectral library matching alone, which often covers less than 10% of the total metabolome. *eCOMET* mitigates this limitation by leveraging class-level annotations (from the *CANOPUS* module of *SIRIUS*; Dührkop *et al*., 2021), which provide hierarchical chemical identities for a much larger proportion of the dataset. Analogous to Gene Ontology (GO) enrichment analysis and gene set enrichment analysis, *eCOMET* performs Chemical Class Overrepresentation Analysis (cORA) and metabolite set enrichment analysis (MSEA; Box 1). This allows researchers to determine which biosynthetic classes are overrepresented among their target features without exact structural annotations (Fig. 1d and 3a).

**Module 3. Quantifying chemical alpha diversity** Another common goal in ecometabolomic studies is the direct comparison of observed chemical diversity between samples and sample groups. eCOMET facilitates such approaches by applying ecological diversity concepts (Fig. 1e and 3b) providing functions and plots for visualizing chemical diversity at the sample or group level and allowing users to optionally filter by biosynthetic pathway annotation. We implemented a Hill number framework (Chao *et al*., 2014; Petrén *et al*., 2023) which unifies richness and evenness into a single metric controlled by the *q* parameter. One way to compute chemical diversity is to use feature quantification data alone, which is particularly useful for intraspecific studies where samples share most features and where diversity is mostly driven by evenness. In interspecific studies, however, treating every unique feature as equally distinct from every other feature can artificially inflate diversity. The functional Hill number approach implemented in *ChemoDiv* takes into account structural similarity and is adopted in eCOMET. This approach ensures that a sample containing structurally distinct compounds (e.g., representing different biosynthetic pathways) is recognized as more diverse than one containing closely related compounds.

**Module 4. Assessing chemical dissimilarity as beta diversity** Lastly, eCOMET provides multiple ways to calculate the chemical distance between samples (Fig. 1f and 3b). Consistent with our diversity framework, users can choose between chemically naïve distances (e.g., Jaccard or Bray–Curtis) and structurally informed distances. The latter include the CSCS (Sedio *et al*., 2017) method and UniFrac distances (Junker, 2018), which treat spectral similarity as a proxy for “phylogenetic” distance between features. These metrics allow for a more nuanced understanding of chemical niche space and community assembly.

In combination, these four modules can be configured into diverse analytical pipelines tailored to specific research designs. In the following sections, we demonstrate the utility of eCOMET through two distinct case studies: (1) a controlled treatment-based experiment on plant-herbivore induction, and (2) a large-scale observatory field study exploring metabolomic shifts across environmental gradients. These two case studies also serve as educational material for training new ecometabolomics practitioners. To facilitate this goal, all data and code are available as vignette tutorials within the package and hosted on the package webpage (https://github.com/Phytoecia/eCOMET).

### Fig. 4. Case 1: Herbivore-induced metabolome showing specific metabolism of Arabidopsis thaliana

To demonstrate how eCOMET can be used to identify features associated with a specific treatment, using Differential Accumulation (DA) analysis, we used a publicly available LC-MS/MS dataset of *Arabidopsis thaliana* leaf tissue attacked by *Spodoptera litura* larvae and *Lipaphis pseudobrassicae*, and a control without herbivores (Fig. 4a). The *mzmine*-processed feature quantification table and SIRIUS-based class annotations are provided in Supplementary Data and are installed directly within the eCOMET package. The overall metabolome was visualized using a heatmap (Fig. 4b). Next, we conducted PCA and plotted the results to observe the distribution of the samples in terms of metabolomes; the three treatment groups were well separated (Fig. 4c). We then performed DA analysis by comparing herbivore-free control groups and two groups attacked by *S. litura* and *L. pseudobrassicae* each. By setting the threshold for DAM to adjusted *p* < 0.1 and |log_2_FC| > log_2_1.5, we found 42 DAMs were increased by both herbivores (Fig. 4d). We then applied cORA on both increased and decreased features to investigate enriched metabolic classes in the DAMs (Fig. 4e). In NPC class-level ontology, the *S. litura*-induced metabolome was enriched with glucosinolates, while the *L. pseudobrassicae*-induced metabolome was enriched with amino acids and monosaccharides. These results illustrate how eCOMET can be used to identify ecologically dynamic molecular features and their metabolic classes.

**Figure 4.**
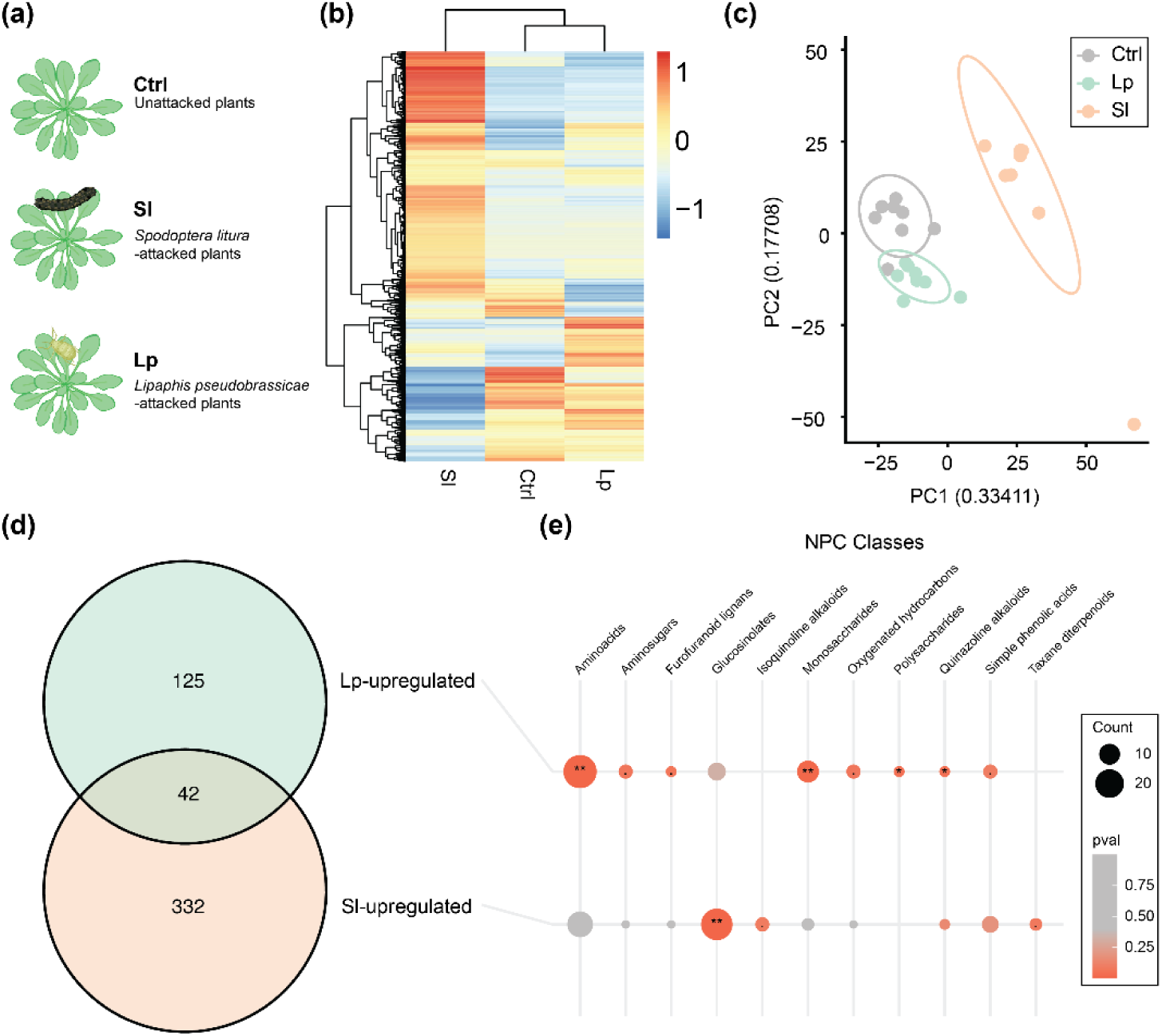
Case 1: Treatment-based metabolomic study. (a) Experimental design of herbivory-induced plant metabolomics. *Arabidopsis thaliana* plants attacked by *Spodoptera litura* and *Lipaphis pseudobrassicae* were sampled and analyzed by LC-MS. (b) Heatmap of all features visualizing Z-socred feature abundance across three groups. (c) PCA plot of samles. (d) Venn diagram of DAMs induced by *S. litura* and *L. pseudobrassicae*. (e) Chemical class overrepresentation analysis of DAMs induced by *S. litura* and *L. pseudobrassicae*.

### Fig. 5. Case 2: Metabolome diversity and dissimilarity among co-occurring tropical tree species

To demonstrate the interspecific ecometabolomics workflow, we used a publicly available LC-MS/MS dataset of foliar metabolome data from tropical tree species sampled in the tropical Andes (Henderson *et al*., 2025; Chadwick *et al*., 2026). This case study demonstrates how to quantify chemical richness and disparity in a structurally weighted or unweighted manner. We first assessed chemical alpha diversity at the sample and species levels using *GetAlphaDiversity*, calculating both feature richness (Fig. 5a) and rarefied richness accumulation curves to account for differences in replicate number across species (Fig. 5b). Chemical richness estimates varied considerably across the 10 focal tree species, and accumulation curves indicated that the numbers of replicates were high enough to approach saturation in chemical sampling for most species.

**Figure 5.**
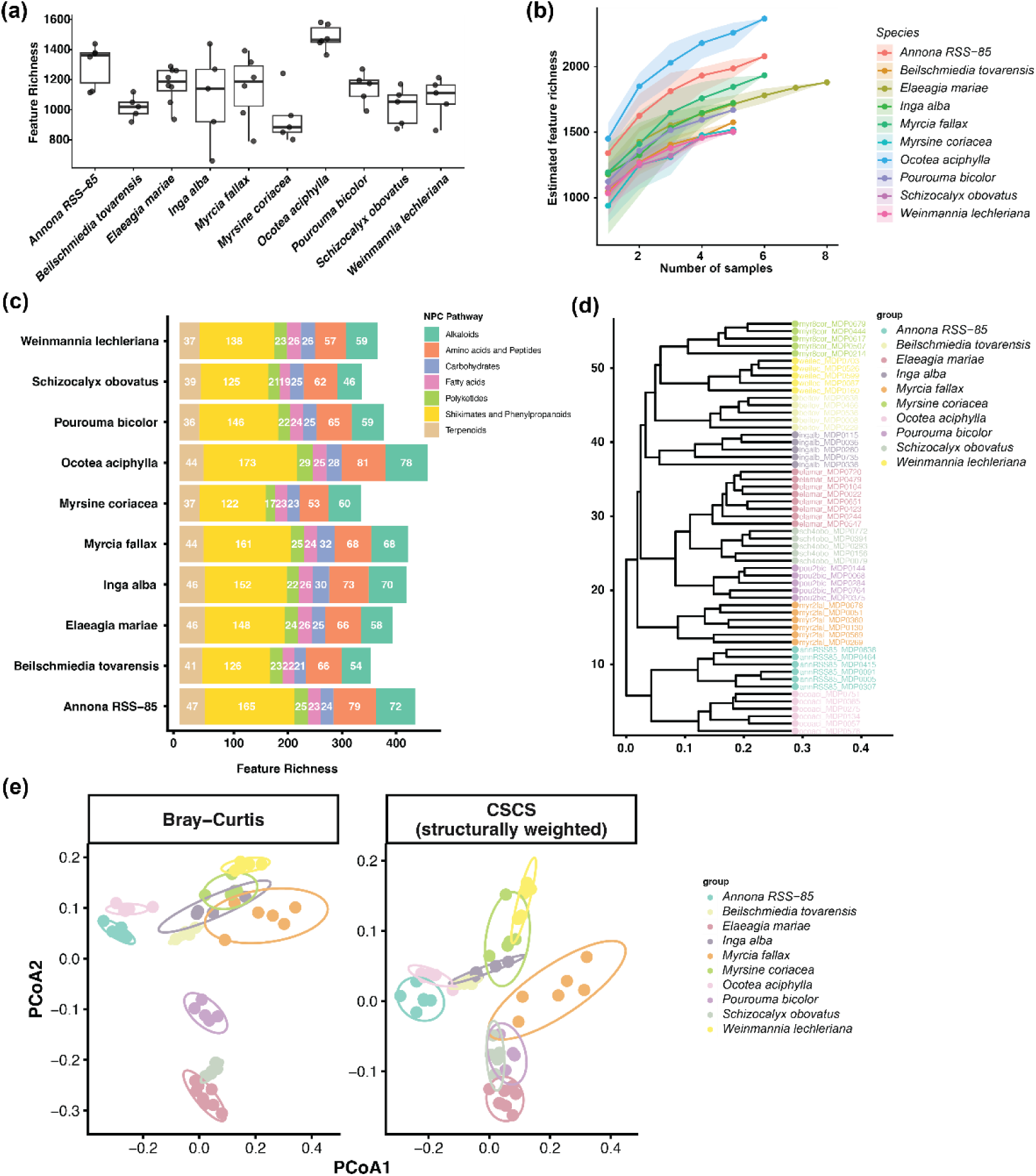
Case 2: Metabolome diversity and dissimilarity among co-occurring tropical tree species. (a) Richness of metabolic features detected from each species. (b) Rarified richness accumulation curves of each species. (c) Output of *PlotNPCStackedBar()* visualizes richness of each NPC pathway annotated by CANOPUS. (d) Hierarchical clustering of samples based on Bray-Curtis distances. (e) PCoA plots of samples based on Bray-Curtis dissimilarity and CSCS.

We then incorporated SIRIUS-based CANOPUS class annotations using *AddSiriusAnnot* and summarized metabolic class composition across species using *PlotNPCStackedBar*. NPC class-level summaries revealed how compound class profiles vary among tree species (Fig. 5c). To characterize beta diversity, we first applied feature-based Bray–Curtis hierarchical clustering, which produced strong species-level clusters, reflecting the limited exact feature overlap among species in this interspecific context (Fig. 5d). Because feature-overlap metrics can overestimate chemical dissimilarity when samples contain structurally related but non-identical compounds, we next introduced comparisons that took into account chemical structure using DreaMS-derived pairwise spectral dissimilarities added via *AddChemDist*. Beta diversity was recalculated using both CSCS and Generalized UniFrac, which incorporate chemical relationships among unshared features. Structural relatedness can be accounted for during clustering by using PCoA on the CSCS distance matrix (Fig. 5e). Finally, functional Hill number weighted by DreaMS spectral dissimilarity was calculated at the sample level, revealing that species with similar raw feature richness could differ substantially in the structural breadth of their metabolomes. These results illustrate how *eCOMET*’s interspecific workflow allows not only feature richness but also chemical diversity estimations that take into account structural similarity, by incorporating ion identity molecular networking (Schmid *et al*., 2021) and therefore providing a more biologically realistic framework for comparing the foliar metabolomes of co-occurring tropical tree species.

### Data sharing

The *mmo* object not only functions as a centralized data format for statistical analysis and visualization, but can also serve as a sharable data format. Exporting and importing *mmo* objects are supported, enabling reproducible analyses and archiving.

### Community Contributions to eCOMET Package

*eCOMET* is an open-source project and the authors welcome contributions from the community, especially those that advance our two main goals: 1) facilitate standardization and reproducibility in ecometabolomics post-processing and visualization, and 2) provide better educational materials to teach ecometabolomics. We encourage ecologists, evolutionary biologists, and analytical chemists to contribute by reporting bugs, requesting new features, or submitting new analysis functions and tutorial vignettes through the project’s GitHub repository (https://github.com/Phytoecia/eCOMET). Detailed contribution guidelines, including code conventions, documentation standards, and dependency policies, are provided on the package website. Feature requests and open-ended discussion about new directions are particularly welcome from users working with study designs or metabolomics workflows not yet represented in the current tutorials.

### Box 1. Chemical term enrichment analyses

Depending on the goals of the study, the proper way to investigate enriched chemical terms can be either chemical class overrepresentation analysis (cORA) or metabolite set enrichment analysis (MSEA). In cases where a set of features of interest can be defined (e.g., DAMs induced by herbivory), cORA is appropriate. cORA tests whether any chemical class is over-represented in this set relative to its frequency in the full metabolome. Specifically, given the number of features belonging to each chemical class in both the set of interest and the full metabolome, a Fisher’s exact test evaluates the probability of observing the class composition of the set under the null hypothesis of random sampling from the full metabolome. In contrast, many ecological studies use correlations of features. When scores can be assigned to individual features (e.g., correlation coefficient with elevation), MSEA can test whether the chemical classes are enriched based on score values. These two approaches to investigating enriched chemical classes are complementary and enable diverse study designs.

## Acknowledgements

We thank C. R. Hawk, E. G. Thurau, M. Volf, T. W. N. Walker, D. Salazar, M. Wang, and E. Gaquerel for helpful discussions. This work was supported by National Research Foundation of Korea (RS-2021-NR061394, RS-2023-00301976, RS-2026-25475700, RS-2024-00434162, and NRF-2022R1A4A3024451) and New Faculty Startup Fund from Seoul National University to MSC and YJ. This work was also supported by US National Science Foundation grant DEB 2240430, and University of Texas at Austin Stengl-Wyer grant SWG-22-01, and Stengl-Wyer Scholars postdoctoral fellowships to DLF and GJD.

## Author Contributions

All authors conceived the study and wrote the manuscript. MSC, DLF, and GJD created the R package with input from BES and YJ.

## Data availability

All data and scripts are available in our GitHub repository (https://github.com/Phytoecia/eCOMET).

## Supporting information

Supporting information contains one figure describing a detailed workflow implemented in *eCOMET* and one table describing ecometabolomic questions *eCOMET* aims to address.

