## Supplementary material for "eCOMET: An R package for evaluating metabolic diversity and enrichment from LC-MS/MS data to test ecological hypotheses from individuals to ecosystems": Figure S1

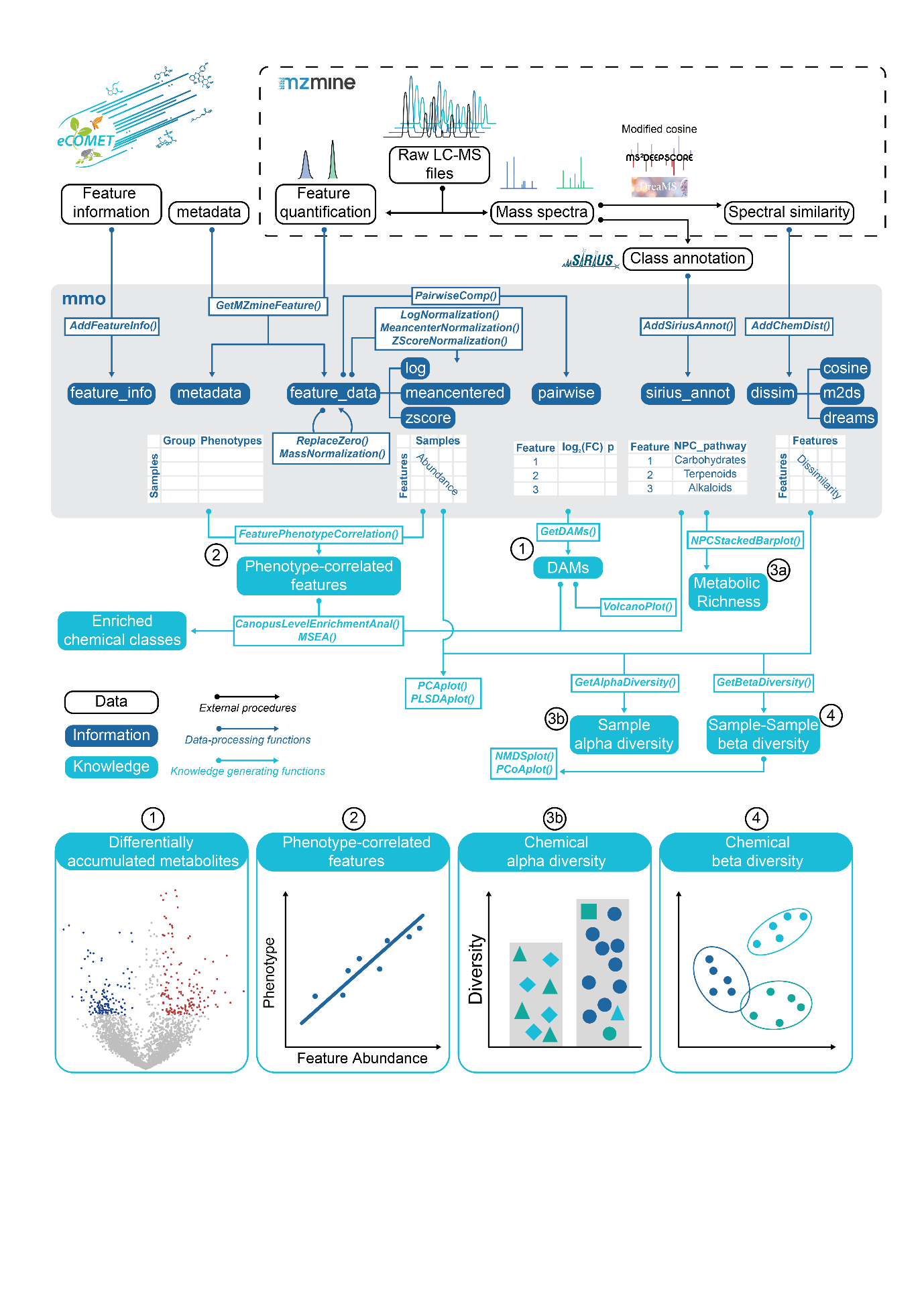

**Suporting information Figure S1.** Detailed structure of eCOMET from data integration to analytic modules.
